# Non-transmissibility of Pepper whitefly borne vein yellows virus (PeWBVYV) by the Mediterranean species of *Bemisia tabaci* and identification of a candidate insect receptor protein for poleroviruses

**DOI:** 10.1101/2020.11.12.380139

**Authors:** Saptarshi Ghosh, Vinicius Henrique Bello, Murad Ghanim

## Abstract

Recent reports of transmission of poleroviruses by whiteflies is indicative of evolution of new virus-vector relationships. Pepper whitefly-borne vein yellows virus (PeWBVYV), was the first report of a polerovirus infecting pepper in Israel which was transmitted by whiteflies (MEAM1) and not aphids. This study reports the inability of the Mediterranean species (MED, Q biotype) of *B. tabaci* to transmit PeWBVYV. However we show that non-transmission of PeWBVYV by MED is not due to the lack of interaction with the GroEL protein of the *Hamiltonella* symbiont. Although not transmitted by MED, PeWBVYV was detected in its hemolymph, indicating its translocation across the MED midgut barrier. The aphid transmitted *Pepper vein yellows virus 2* (PeVYV-2) was also detected in the hemolymph of MEAM1 whiteflies but PeWBVYV could not be detected in the aphid hemolymph. Interestingly, relative amounts of PeWBVYV in the hemolymph of the, MED was much lower than in hemolymph of MEAM1 whiteflies. We also identified a candidate receptor protein, complement component 1Q sub-complement binding protein (C1QBP) which interacts with the capsid proteins of PeWBVYV and PeVYV-2 but not with the whitefly transmitted *Tomato yellow leaf curl virus* by a yeast two-hybrid approach using the minor capsid protein (RTD) as bait to screen for interacting proteins against the whitefly cDNA library. C1QBP, is a known receptor of bacterial and viral pathogens but this is the first report of its interaction with a plant virus.

## Introduction-

The majority of plant viruses require an insect vector for transmission onto new hosts. However, insect transmission of plant viruses is highly specific, ensured by stringent interactions between the viruses and their insect vectors (1). Virus genetic elements determining insect transmission are thus conserved to maintain insect transmissibility, which impedes virus evolution and also restricts emergence of new virus-vector interactions (1, 2). Recent reports of non-canonical transmission of recombinant poleroviruses by whiteflies (*Bemisia tabaci*) and not aphids despite high genome identity with other aphid-transmitted poleroviruses (3, 4) although mark an evolutionary leap and perplexes our understanding of virus-insect vector interactions. *B. tabaci*, a cryptic species complex transmits over 300 plant viruses from the genera *Crinivirus*, *Ipomovirus*, *Torradovirus* and *Carlavirus* semi-persistently and from *Begomovirus* in a persistent mode (5). The persistently transmitted begomoviruses although much different from poleroviruses, yet are similar with respect to the transmission route and tissue specificity (6), which might be a contributory factor towards emergence of polerovirus-whitefly relationships.

The circulative pathway of the virions after ingestion by its specific aphid or whitefly vector involves recognition and adsorption to the membrane of the insect hindgut or posterior midgut for aphids (7, 8) while the filter chamber or anterior midgut for whiteflies (9, 10); internalization and translocation across the gut epithelial cells into the hemocoel by receptor mediated endocytosis (11–14); streaming in the hemocoel to reach the accessory salivary glands for aphids (15) and primary glands for whiteflies (9); and transportation across the glands to the salivary duct (6, 16) before egestion onto the next host plants. Throughout this process, the insect midgut (7, 17) and the salivary glands (18–20) of aphids and whiteflies serve as anatomical barriers determining the fate of the virions. Transmissibility of circulative viruses by aphids or whitefly is influenced simultaneously by virus and insect factors. Capsid proteins of both luteoviruses (21, 22) and begomoviruses (20, 23) are the only known virus factors determining its transmission by aphids and whiteflies, respectively. Conserved domain features of the capsid proteins of poleroviruses and begomoviruses are indicative of its involvement in interaction and transport across the tissue barriers within aphids (6) and whiteflies (20, 24), respectively.

The co-existence of two poleroviruses infecting pepper crops in Israel, the aphid transmitted Pepper vein yellows virus 2 (PeVYV-2) and the whitefly transmitted Pepper whitefly-borne vein yellows virus (PeWBVYV) is aided by their transmission by two discrete insect vectors (25). The viral genetic element determining the specific transmission of the two viruses by aphids and whiteflies remains an enigma but can be an ideal model to decipher the stringent requirements of specific virus-vector transmission. Aphid transmissibility of poleroviruses is determined by the virus structural proteins which includes a major capsid protein (CP, 22 kDa) and a read-through minor capsid protein (RTD, 55 kDa) (11). Point mutations of conserved regions of the RTD reduced virion transport both across the gut and the accessory salivary glands, reduced virion stability in the hemolymph and subsequent transmission to host plants, indicating its role in the viral movement, stability and transmission (21, 22, 26–28). The major aphid factors that are hypothesized to affect luteovirid transmission are receptors which reside on the gut-hemolymph (6, 18, 29, 30) and the hemolymph-accessory salivary glands barriers (15, 18, 31). Furthermore, it has been shown that the stability of virions in the hemolymph is mediated by interaction of the RTD with a GroEL protein produced by the primary symbiont of aphid *Buchnera* (32).

Interaction with the GroEL chaperone secreted by *Hamiltonella*, a facultative bacterial endosymbiont harbored in the *B. tabaci* also known to increase stability of begomoviruses within the whitefly hemolymph (33, 34). This interaction of the begomovirus capsid protein with the *Hamiltonella* GroEL is also hypothesized to condition Middle East Asia Minor 1 (MEAM1 or B biotype) species of *B. tabaci* as better vectors of the *Tomato yellow leaf curl virus* (TYLCV) than the Mediterranean (MED or Q biotype), which lacks *Hamiltonella* (35). MEAM1 whiteflies have been previously shown to be the vector for PeWBVYV (3). In the current study, we investigated the transmission efficiency of PeWBVYV by the MED species of *B. tabaci* and report its non-transmissibility by MED species. To elucidate whether similar to TYLCV, there is a role of the GroEL protein of *Hamiltonella* behind the exclusive transmission of PeWBVYV by MEAM1, we used a yeast two-hybrid (Y2H) assay to compare binding of the RTD proteins of PeWBVYV/PeVYV-2 with the *Hamiltonella* GroEL protein. The Y2H approach was also used to screen the viral minor capsid protein (RTD) against the whitefly cDNA library for identification of complement component 1 Q sub-component binding protein (C1QBP) as a putative receptor of PeWBVYV. Interactions of the aphid transmitted PeVYV-2 and PeWBVYV with the C1QBP of *B. tabaci* and the green peach aphid (*Myzus persicae*) were compared. Moreover, we compared the ability of the PeWBVYV to cross the midguts of MEAM1 and MED to reach the hemolymph.

## Materials and method

### Transmission of PeWBVYV by *B. tabaci* MED species

*B. tabaci* (MED) (~25 whiteflies/plant) reared on cotton plants were given acquisition access to PeWBVYV for 72 hours followed by life-long inoculation access onto pepper plants in three replicates. A control experiment was set up with MEAM1 whiteflies reared on cotton and allowed acquisition access to PeWBVYV for 72 hours with subsequent inoculation to twenty pepper plants. Sample whiteflies collected after acquisition and inoculation access were tested for the presence of PeWBVYV. Total RNA was extracted from top leaves of inoculated pepper plants at 25 days post inoculation (dpi) and screened for infection with PeWBVYV by RT-PCR as described in Ghosh et al. 2019 (3). All experiments and insect colony maintenance were conducted under controlled environment (25°C ± 5, 60% RH, 14hr L: 10hr D).

### Detection and quantification of PeWBVYV/PeVYV-2 from hemolymph of aphids and whitefly-

Hemolymph of whiteflies (MEAM1 and MED) and aphids were extracted to ascertain whether PeVYV-2 and PeWBVYV cross the midguts of the non-transmitting insects, respectively. Non-viruliferous aphids and MEAM1/MED species of *B. tabaci* were allowed 72 hours of acquisition access to both PeVYV-2 and PeWBVYV separately and collected. A sample of whiteflies and aphids were homogenized for RNA extraction to confirm the presence of the viruses acquired. Hemolymph drawn using pulled glass micropipettes from the thorax of five adult whitefly females collected after virus acquisition were pooled in a PCR tube containing 10 μl of sterile water. Similarly, hemolymph droplets oozing out from punctured thorax of two aphid nymphs was collected using the pulled micropipettes and was pooled in PCR tubes containing 10 μl of sterile water. Hemolymph collected was directly used for cDNA synthesis (20 μl reaction volume) with virus specific primers using M-MLV reverse transcriptase (Promega, USA). Presence of PeVYV-2 or PeWBVYV was determined by PCR (20 μl reaction volume) with 9 μl of the cDNA as template using virus specific primers (3).

Relative quantities of PeWBVYV normalized to the whitefly actin was quantified from the hemolymh of MEAM1 and MED whiteflies by qPCR as previously described (25). Nine microliters of cDNA was used as template for the qPCR with two replications for each sample and relative quantities were calculated by deltadelta Ct method.

### Whitefly Library construction-

Approximately 300 viruliferous *B. tabaci* (MEAM1) adults with PeWBVYV were homogenized with liquid nitrogen and 50 μl TRI Reagent (Sigma) in 1.5 ml centrifuge tubes with a micro pestle followed by washing of the micro pestle by additional 550 μl TRI reagent. The homogenized sample was incubated at room temperature for 10 minutes, 150 μl of chloroform was added and centrifuged to separate the aqueous and organic phases. The aqueous phase was mixed with 0.5 volumes of 100 % ethyl alcohol and added to spin columns of the RNeasy mini kit (Qiagen). Total RNA was extracted using the RNeasy kit protocol. The extracted RNA was quantified and assessed for quality by Nanodrop ND1000. First strand cDNA was synthesized from 3 μg of the RNA using the Make your own mate & plate library system (Takara Clontech) with the CDS III adaptor primer as per manufacturer’s instructions with slight modification of extended extension for 20 minutes before addition of the template switching Smart III modified oligo. The first strand cDNA was amplified in 18 thermal cycles by Long distance PCR using the Advantage 2 polymerase mix (Takara Clontech) as per manufacturer’s instructions. An aliquot of the PCR product was analyzed in 1% agarose gel for estimation of fragment range and quality followed by purification of the remaining amplified product using chroma spin + TE 400 columns to select fragments above 400 bp. The purified ds CDNA was co-transformed along with pGADT7-Rec into competent Y187 yeast cells using Yeast maker yeast transformation system 2 (Takara Clontech) and plated on–Leucine SD minimal agar media (Takara Clontech) as per manufacturer’s instructions. 100 μl of a 1/100 dilution of the transformed cell suspension (15 ml) when plated on–Leucine SD Agar media with 72 hours of growth at 30°C yielded 371 colonies with estimated 5.56 million independent cDNA clones in the library. The colonies were harvested in YPDA freezing media and stored in aliquots of 1.4 ml in cryogenic tubes at −80°C.

### Bait constructs for Y2H screening

The minor capsid proteins of PeWBVYV (W-RTD), PeVYV-2 (Y-RTD) were used as baits for screening interacting proteins from the whitefly. The CP-stop codon was removed from the ORFs of the RTDs of both viruses to ensure translation of only RTD, as a single protein. The RTDs were first PCR amplified in two fragments ending and beginning upstream and downstream of the major coat protein stop codon, respectively and then joined by overlap extension PCR with primers with overlapping sequences excluding the stop codon (Table 1). The full length RTDs were amplified using primers containing 5’ overlapping sequences of the pGBKT7 vector and ligated into linearized pGBKT7 (*Eco*RI + *Bam*HI) using Gibson assembly master mix (NEB). The major capsid protein of PeWBVYV (W-CP) was PCR amplified and cloned into pGBKT7 using Gibson assembly as described before. The TYLCV coat protein gene (TYLCV-CP) was PCR amplified from infected plants and cloned into pGBKT7 as described in Morin et al. 2010 (34). All PCR amplifications in this study were done using Q5 High-Fidelity DNA polymerase (NEB). Empty pGBKT7, pGBKT7-p53 and pGBKT7-Laminarin (Takara Clontech) were used as control baits. All bait constructs in pGBKT7 were cloned into the Y2H Gold yeast strain using the Yeast maker yeast transformation system 2 (Takara Clontech).

**Table 1.**
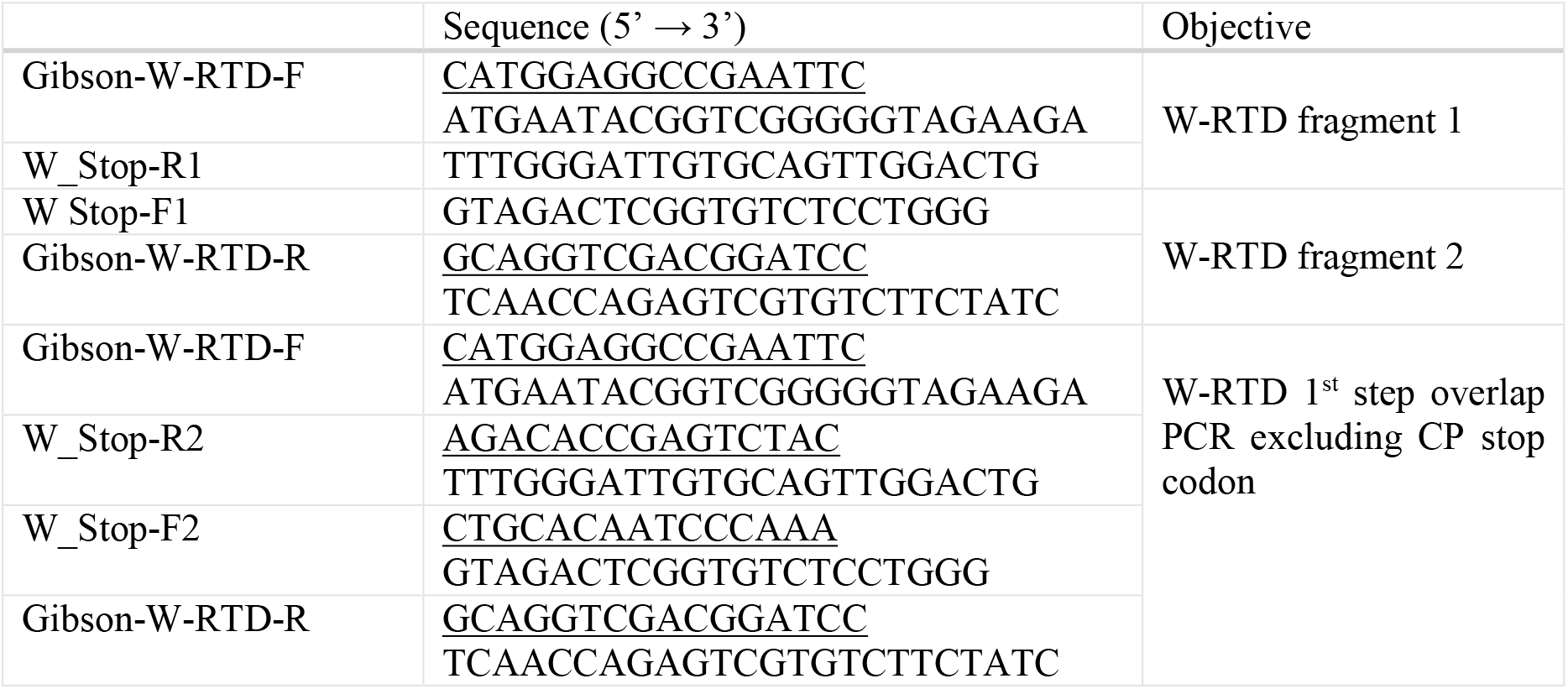
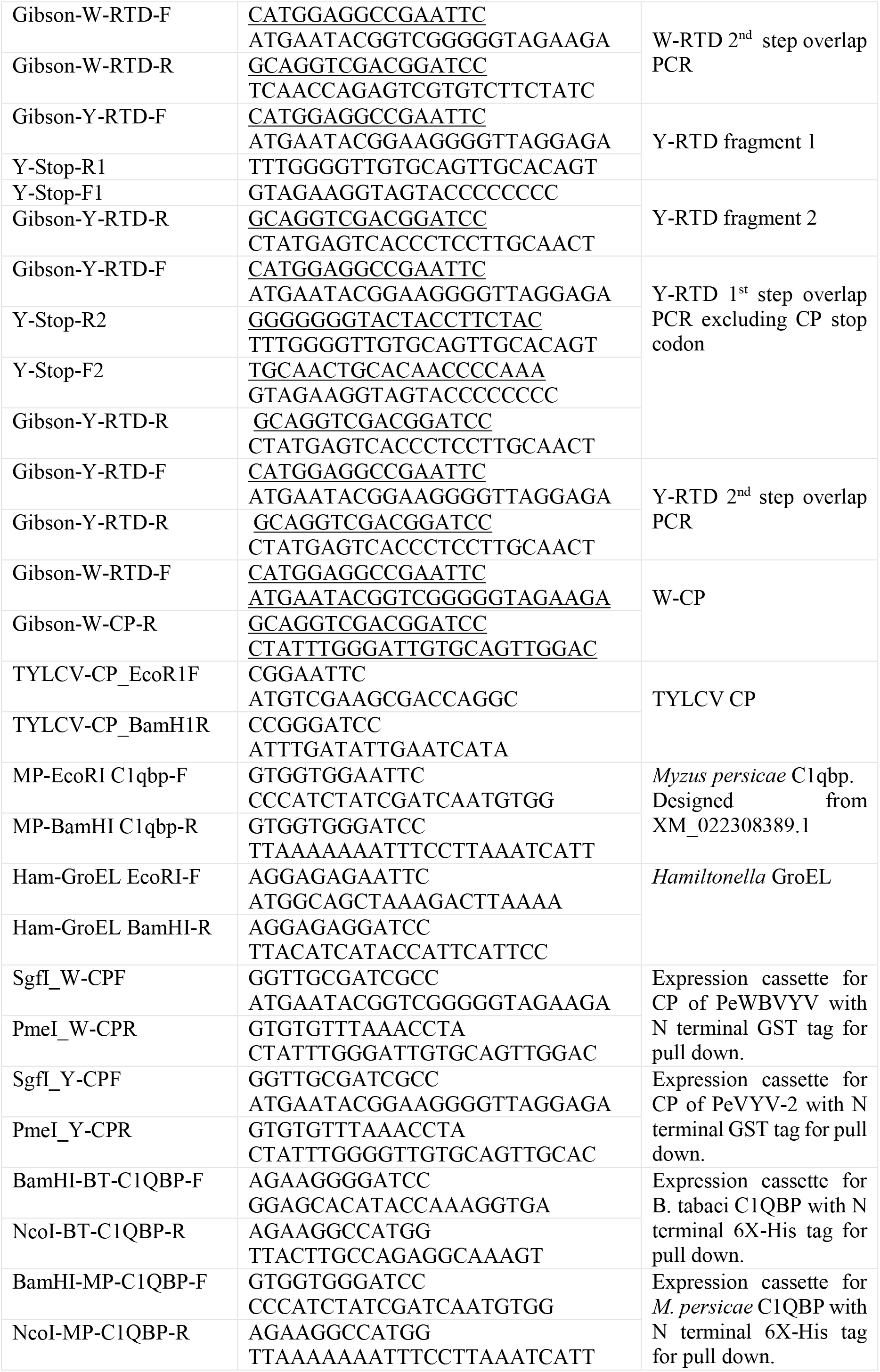
Primer used in this study. The underlined sequences represent the overlapping sequences of the vector used for Gibson assembly.

### Prey constructs for Y2H screening

The GroEL gene of the secondary endosymbiont *Hamiltonella* was used as a prey to screen against the RTD bait proteins. The *Hamiltonella* GroEL was PCR amplified using primers (34) with additional *Eco*RI and BamHI sequences (Table 1) at the 5’ end and cloned into pGADT7 by restriction enzyme cloning using *Eco*RI and *Bam*HI. The C1QBP gene of the peach aphid, *Myzus persicae* (*M. persicae*) was also used as a prey in this study. A partial gene of the *M. persicae* C1QBP protein (XP_022164081.1) was PCR amplified from *M. persicae* cDNA using specific primers (Table 1) and cloned into pGADT7.

Empty pGADT7 and pGADT7-T antigen were used as control preys. All prey constructs in pGADT7 were cloned into the Y187 yeast cells (Takara Clontech).

### Yeast two hybrid assays

The W-RTD and Y-RTD bait constructs were screened against the *Hamiltonella* GroEL prey construct. A single colony (Y2H Gold) of W-RTD and *Hamiltonella* GroEL (Y187) each were inoculated to 500 μl of 2X YPDA broth, allowing mating for 24 hours at 30°C at 200 rpm. 100 μl of the culture was plated onto SD agar minimal media lacking leucine and tryptophan (–LT) and incubated at 30°C for 72 hours. Colonies developed were screened on SD minimal agar media lacking adenine, leucine, tryptophan and histidine (–ALTH). Colonies from–LT media were also screened on less stringent minimal media lacking leucine, tryptophan and histidine (–LTH). Y-RTD interactions with *Hamiltonella* GroEL was also screened as described above.

The W-RTD bait construct was screened against the *B. tabaci* cDNA library prey constructs as described in the Matchmaker Gold yeast two-hybrid system user manual (Takara Clontech). Re-suspended cells after mating were plated on–ALTH media and incubated at 30°C for 8 to 14 days. Colonies developed were streaked on to–ALTH media with X-alpha-Gal (–ALTH/X) and screened for development of blue color. Blue colored colonies were further screened on–ALTH/X supplemented with Aureobasidin A (40 μg/ml) (–ALTH/X/A). The length of the prey sequence was analyzed from the blue colonies by colony PCR using Matchmaker AD LD Insert screening primers. Also, the presence of W-RTD bait was verified from the blue colonies by colony PCR using T7 promoter and W-RTD specific reverse primers. Colonies developed on–ALTH/X/A were further inoculated on–LT minimal broth and the plasmids were isolated (http://mapageweb.umontreal.ca/rokeach/protocoles/extraction_EN.html). The pGADT7-whitefly cDNA construct was recovered by transformation into *E. coli* and selection on LB agar with ampicillin. The pGADT7-whitefly cDNA plasmid after isolation was Sanger sequenced from both directions. The isolated pGADT7-whitefly cDNA plasmids were re-transformed into Y-187 strains and screened against the W-RTD, Y-RTD, W-CP, TYLCV-CP and pGBKT7 without insert bait constructs each by one-to-one mating assays.

C1QBP from *M. persicae* was PCR amplified using specific primers (Table 1) and screened against the PeWBVYV and PeVYV-2 bait proteins.

### Sequence analysis, annotation, homology modelling and phylogeny-

The open reading frames of the prey nucleotide sequences obtained from the Y2H assays were checked to be in frame with the GAL4 activation domain and were annotated using BLASTp algorithm and the pfam (36), CDD (37) databases. Amino acid sequences of C1QBP from hemipteran insect vectors were aligned using the MEGA 7.0 software (38). The C1QBP gene sequence of the MED *B. tabaci* was obtained by BLAST against the available genome at Genbank (GCA_003994315.1) and was also amplified by RT-PCR from total RNA extracted from MED whiteflies. Phylogenetic analysis of aligned amino acid sequences were analyzed using the MrBayes on XSEDE tool with fixed LG+G substitution model from the CIPRES gateway (39). The phylogenetic analysis was run for 1 million generations with sampling every 1000 generations. Homology modelling of the C1QBP proteins of *B. tabaci* and *M. persicae* were done using the SWISS-MODEL with the P32 mitochondrial matrix protein of humans (SMTL ID: 1p32.1) as template. The predicted models were aligned using PyMOL to compare their 3D structures.

### Pull down of whitefly/aphid C1QBP by coat protein of PeWBVYV/PeVYV-2-

The CP gene of PeWBVYV and PeVYV-2 were inserted into pFN2A vector (Promega, USA) to express the CP as fused proteins with N terminal GST tag. pFN2A vector with a GFP insert was used as GST control. Similarly, partial C1QBP from *B. tabaci* and *M. persicae* were cloned into pRSETA vector to express as fused proteins with N terminal 6xHis tag. The constructs were transformed into BL21 (DE3) (New England Biolabs, UK) strain of *E. coli* and protein expression was induced using 0.4 mM IPTG at 37°C for 4 hours. The GST-CP and GST-GFP fusion proteins from soluble fractions of the bacterial lysate were immobilized onto MagneGST™ particles (Promega, USA) and were used as baits to pull down C1QBP proteins from soluble fractions of the bacterial lysates expressing the 6xHis-C1QBP fused proteins as per manufacturer’s protocol. The prey proteins after elution with boiling in 1x SDS buffer were electrophoresed in 10% SDS-PAGE gel and detected by western blotting using monoclonal anti-poly histidine antibody (Sigma Aldrich, Israel).

### Localization of PeWBVYV inside whitefly tissue by fluorescent in situ hybridization (FISH)

Midguts and salivary glands from viruliferous and non-viruliferous whiteflies (MEAM1) were dissected in citrate buffer saline (CBS, pH 6.5) and fixed with 4% paraformaldehyde for 30 minutes. The midguts and salivary glands were washed thrice with CBS and incubated overnight with hybridization buffer [1 mM EDTA, 40 mM PIPES (pH 6.4), 400 mM NaCl, 40% formamide] containing Cy3-labelled probe (100 mM) specific to PeWBVYV (3). The dissected organs were washed thrice with CBS and mounted on slides with hybridization buffer containing DAPI (10 μg/ml) and viewed under a confocal laser scanning microscope.

## Results

### PeWBVYV is not transmissible by MED species of *B. tabaci*

PeWBVYV was not transmitted to any of the pepper plants (0 out of 45) inoculated with viruliferous MED species of *B. tabaci* (Fig. 1 A). Acquisition of PeWBVYV by the MED whiteflies was confirmed by PCR on whitefly samples collected after acquisition access. In contrast, 90% (18/20) of the pepper plants inoculated with control MEAM1 whiteflies were detected with PeWBVYV (Fig. 1B).

**Figure 1.**
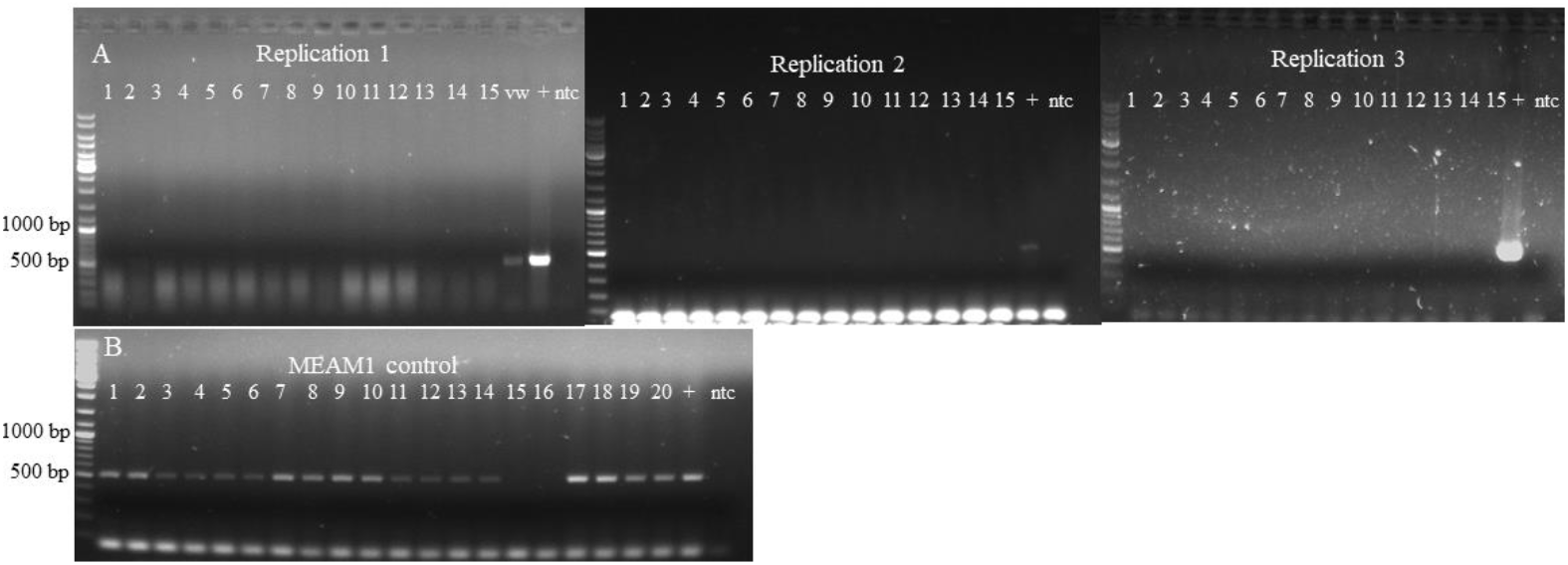
RT-PCR detection of PeWBVY in bell-pepper plants inoculated with viruliferous MED (A) and MEAM1 control (B) whiteflies. vw denote MED whiteflies after acquisition access to PeWBVYV and (+) denote PeWBVYV infected plant used for acquisition.

### The RTD-proteins of PeWBVYV and PeVYV-2 do not interact with *Hamiltonella* GroEL

The PeWBVYV and PeVYV-2 RTD proteins, devoid of the stop codon of the major capsid protein were expressed as fusion proteins with the GAL 4 DNA binding domain (BD) that bind to unrelated promoters of four reporter genes (HIS3, ADE2, AUR1-C and MEL1) in the Y2H Gold yeast strain. The GroEL protein of *Hamiltonella* symbiont of the *B. tabaci* (MEAM1) was expressed fused with the GAL4 Activation domain (AD) in the Y187 yeast strain. Diploid yeast cells expressing the two fused proteins DNA BD-W/Y-RTDs and the AD-*Hamiltonella* GroEL (Figure 2A) failed to grow in both the less stringent–LTH and the more stringent–ALTH media (Figures 2 B). Interactions between TYLCV-CP and *Hamiltonella* GroEL was tested as a positive control and diploid yeast cells expressing the fused proteins DNA BD-TYLCV-CP and AD-*Hamiltonella* GroEL developed in–ALTH media with development of blue color (Figure 2C). This indicates that the RTD proteins of PeWBVYV and PeVYV-2 do not interact with the GroEL protein of the *Hamiltonella* symbiont of *B. tabaci*.

**Figure 2.**
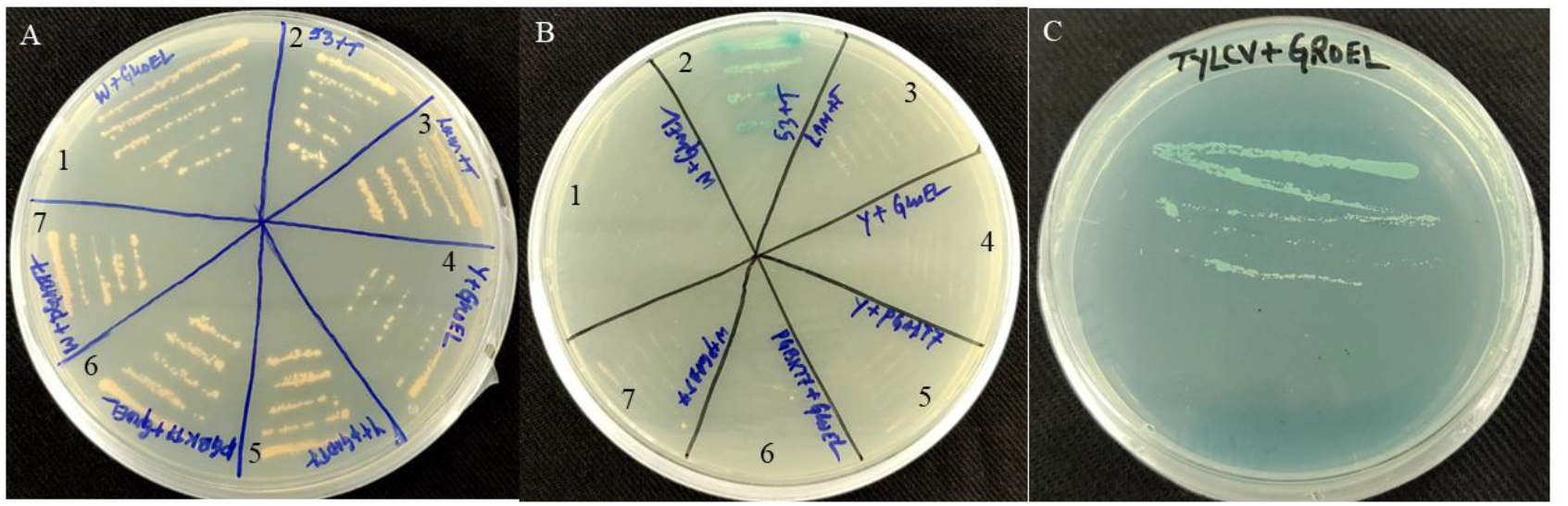
Y2H assay of W/Y-RTD against *Hamiltonella* GroEL. Diploid yeast colonies containing bait + prey constructs of W-RTD + GroEL (1), p53 + T antigen (2), Laminarin + T antigen (3), Y-RTD + GroEL (4), Y-RTD+pGADT7 (5), pGBKT7+GroEL (6) and W-RTD + pGADT7 on–LT (A) and–ALTH/X media (B). Interactions between TYLCV-CP (bait) and *Hamiltonella* GroEL (prey) (C) used as positive control for this assay.

### Detection and quantification of PeWBVYV/PeVYV-2 from hemolymph of aphids and whitefly-

To know whether PeVYV-2 (transmitted by aphids) crosses the midgut barrier of the whiteflies, we screened the hemolymph of MEAM1 whiteflies by RT-PCR after 72 hours of acquisition access to PeVYV-2. PeVYV-2 could be detected in the hemolymph of all the 10 samples tested (Fig. 3 A, B). Hemolymph extracted from MEAM1 whiteflies with acquisition access to PeWBVYV was used as positive control (Fig. 3 C). Interestingly, PeWBVYV was also detected from the extracted hemolymphs of MED whiteflies with acquisition access to PeWBVYV (Fig. 3 D). Our results imply that both PeVYV-2 and PeWBVYV can cross the midgut barrier of the non-transmitting MEAM1 and MED whiteflies, respectively. In contrast, we could not detect the whitefly transmitted PeWBVYV in the hemolymph of aphids allowed 72 hour acquisition access to PeWBVY (Fig. 3 E, F). PeVYV-2 was detected in the hemolymph of the control aphids with acquisition access to PeVYV-2 (Fig. 3 G).

**Figure 3.**
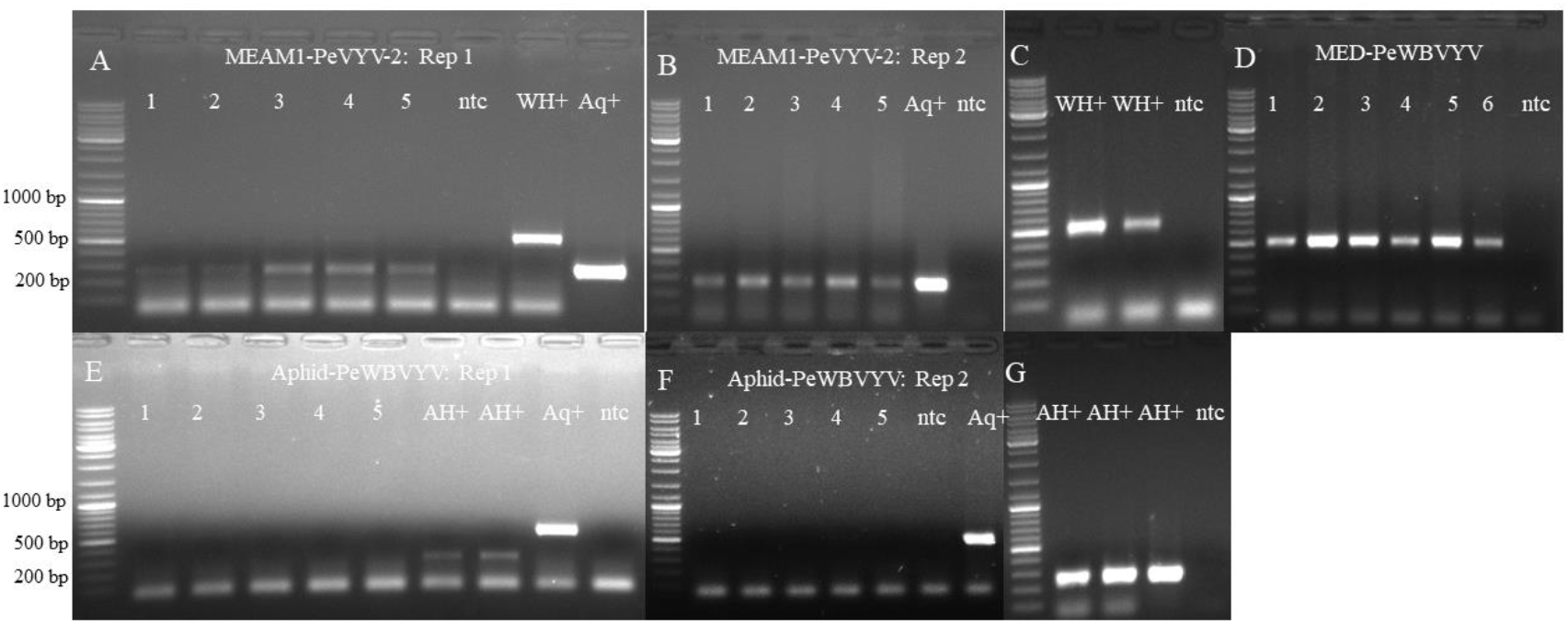
RT-PCR detection of PeVYV-2 from pooled hemolymph samples extracted from five MEAM1 whiteflies after 72 hours of acquisition access to PeVYV-2 (A-B) with hemolymph samples from five MEAM1 whiteflies (WH+) with 72 hours acquisition access to PeWBVYV as positive control (C). RT-PCR detection of PeWBVYV from pooled hemolymph samples extracted from five MED whiteflies after 72 hours of acquisition access to PeWBVYV (D). RT-PCR detection of PeWBVYV from hemolymph samples extracted from two aphid nymphs after 72 hours of acquisition access to PeWBVYV (E-F) with hemolymph samples from two aphid nymphs (AH+) with 72 hours acquisition access to PeVYV-2 as positive control (G). The acquisition of PeVYV-2 and PeWBVYV by MEAM1 whiteflies and aphid, respectively were confirmed from pooled samples collected after acquisition (Aq+).

Interestingly, relative amounts of PeWBVYV (normalized to the actin gene of the whitefly) acquired after 72 hours of acquisition access although was indifferent (F=1.73, P = 0.2) in MEAM1 and MED whiteflies (whole insects), yet significantly higher titers (>25 fold) of PeWBVY was detected in the hemolymph of MEAM1 than in the non-transmitting MED whiteflies (Fig. 4). Similarly, PeVYV-2 although was shown in this study to cross the midgut barrier of the MEAM1 whitefly, however was detected in very low amounts (<1000 fold) in the hemolymph compared to PeWBVYV in MEAM1 (Fig. 4).

**Figure 4:**
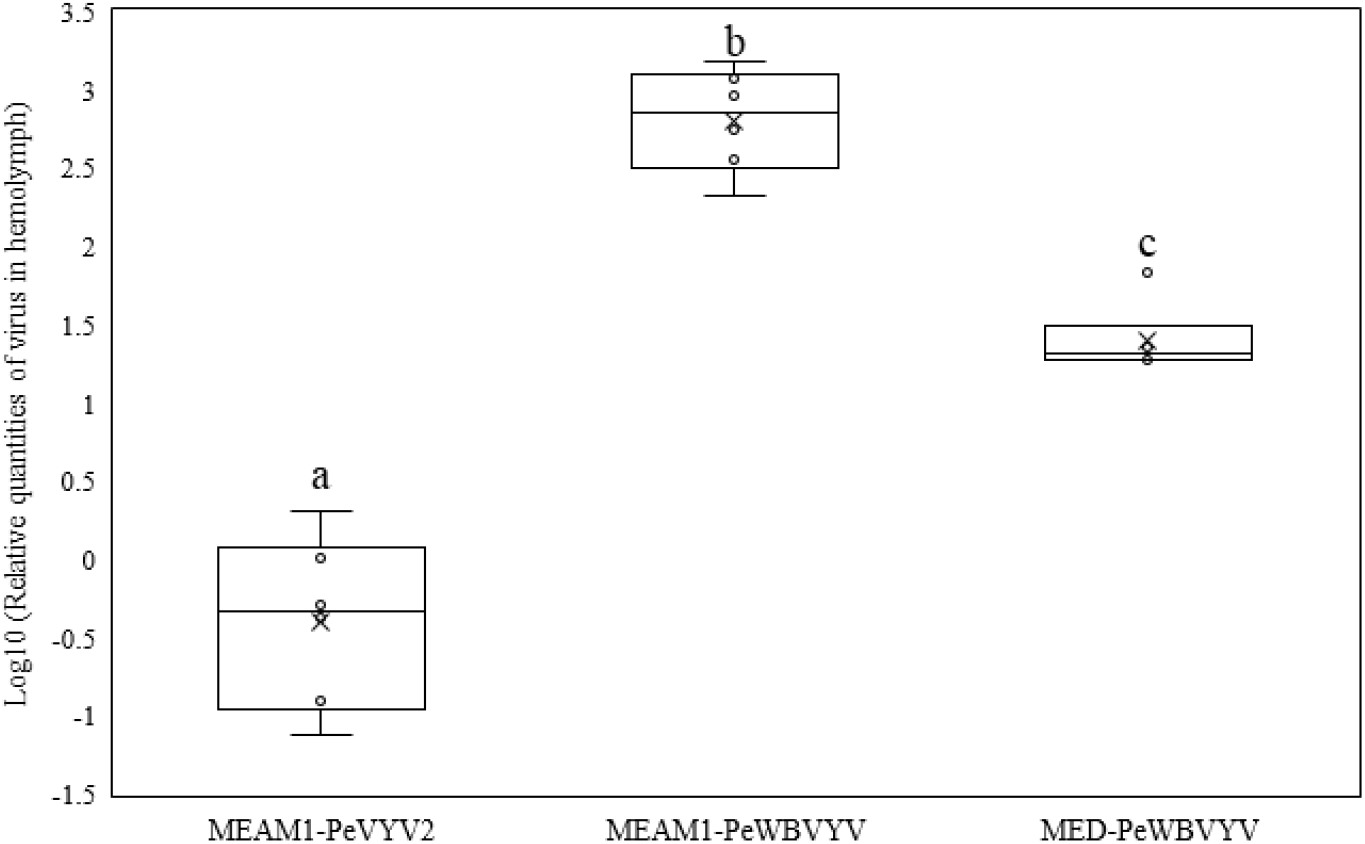
Relative quantification of PeVYV-2 and PeWBVYV in the hemolymph of MEAM1 whitefly and PeWBVYV in MED hemolymph.

### The C1QBP protein of the aphid and whitefly interact with the capsid proteins of PeWBVYV and PeVYV-2

Screening of the fused DNA BD-W-RTD bait construct against the *B. tabaci* cDNA library prey constructs resulted in seven colonies on the–ALTH most stringent media, out of which two colonies were identified to contain the complement component 1 Q sub-component binding protein (C1QBP) gene sequence of the *B. tabaci* as the prey. BLASTp analysis of this prey sequence confirmed its identity with 100% sequence identity with C1QBP protein (XP_018897713.1) from *B. tabaci* (MEAM1). The C1QBP prey transcript identified by the Y2H assay was partial, encoding from the 73^rd^ amino acid of the full C1QBP protein of *B. tabaci* (Fig. 5 A). The isolated C1QBP prey plasmid was retransformed into Y187 yeast strain and was screened against the W-RTD, Y-RTD, W-CP, TYLCV-CP and pGBKT7 constructs on–ALTH/X/A media. Partial C1QBP gene of *M. persicae* (Fig. 5 A) was also cloned into pGADT7 (prey vector) and screened against the baits W/Y-RTDs on -ALTH/X/A media. The C1QBP from *B. tabaci* interacted both with the minor capsid proteins of both PeWBVYV (W-RTD) and PeVYV-2 (Y-RTD) and also with the major capsid protein of PeWBVYV (W-CP) (Figures 5 A) indicating the major capsid protein as the interacting protein and not the minor capsid protein (RTD minus the CP). However, no observable interaction of the whitefly C1QBP could be detected with either the coat protein of the TYLCV (TYLCV-CP) or the empty bait vector (pGBKT7) indicating its specific interaction with pepper infecting poleroviruses. We also report interaction between the coat proteins of the pepper infecting poleroviruses used as bait in this study with the aphid C1QBP. Diploid yeast colonies containing either W-RTD or Y-RTD as the bait together with the *M. persicae* C1QBP prey construct developed in–ALTH/X/A media indicating interaction (Figure 5 B). Interestingly, the interaction of the *M. persicae* C1QBP was stronger with Y-RTD than that of W-RTD as indicated by the development of the blue color (Figure 5 B). Interactions between the C1QBP protein of whiteflies/aphids and the coat proteins of PeWBVYV and PeVYV-2 was further confirmed by *in vitro* pull down assays using the CPs as baits. The results were similar to the Y2H assays wherein the whitefly C1QBP was efficiently pulled down by the coat proteins of both PeWBVYV and PeVYV-2 (Fig. 5 D), and similarly the C1QBP of *M. persicae* interacted with PeVYV-2 and PeWBVYV (Fig. 5 D).

**Figure 5.**
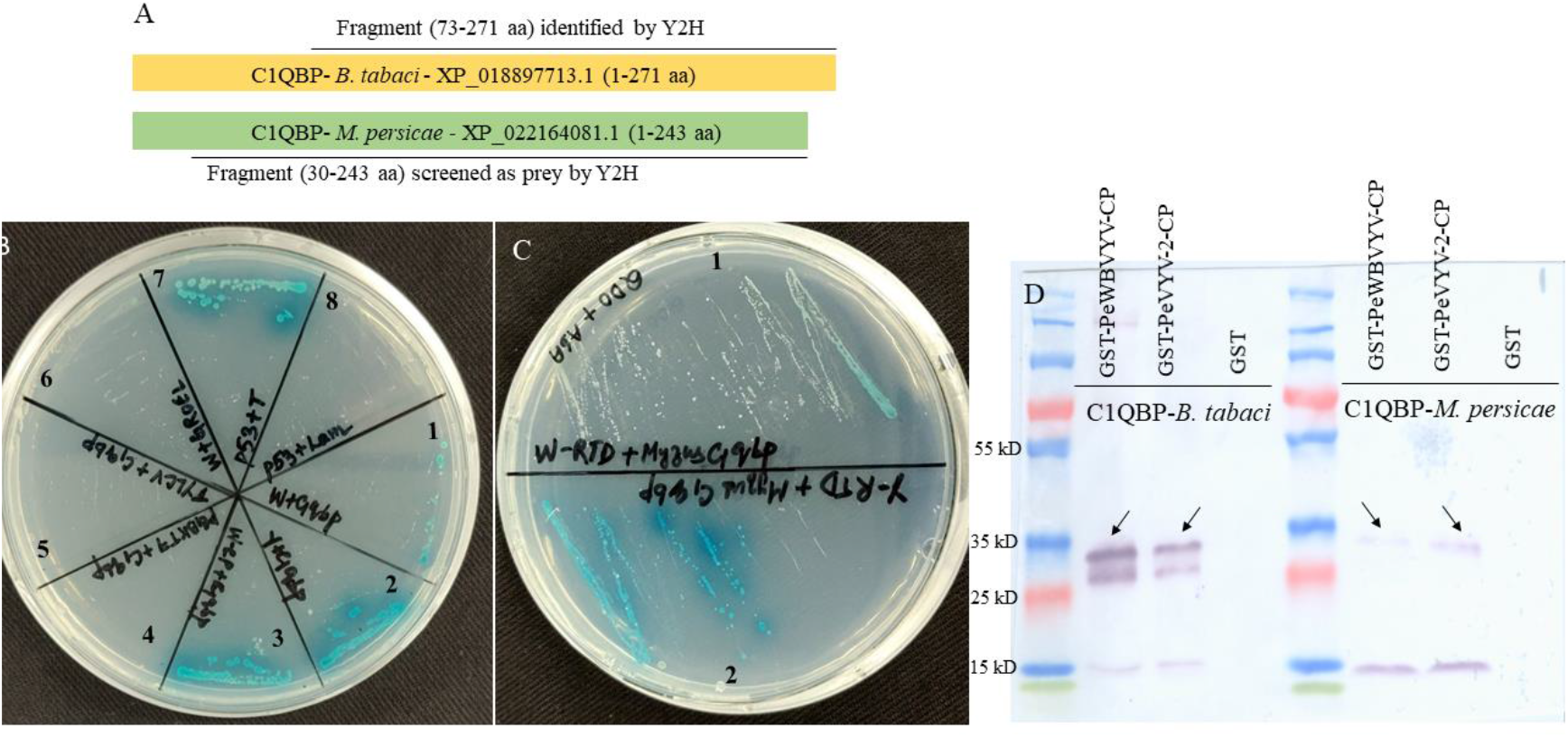
Representation of the C1QBP protein of *B. tabaci* and *M. persicae* and the interacting region used in the Y2H analysis (A). Diploid yeast colonies containing bait + prey constructs of W-RTD + C1QBP of *B. tabaci* (1), Y-RTD + C1QBP of *B. tabaci* (2), W-CP + C1QBP of *B. tabaci* (3), pGBKT7 + C1QBP of *B. tabaci* (4), TYLCV-CP + C1QBP of *B. tabaci* (5), W-RTD + C1QBP of *B. tabaci* (6), p53 + T antigen (7), and Laminarin + T antigen (8) on–ALTH/X/A media (B). Diploid yeast colonies containing bait + prey constructs of W-RTD + C1QBP of *M. persicae* (1) and Y-RTD + C1QBP of *M. persicae* (2) on–ALTH/X/A media (C). Detection of C1QBP of *B. tabaci* and *M. persicae* with N-terminal 6x-his tag after pull down using GST tagged coat proteins (CP) of PeWBVYV and PeVYV-2 as baits by western blot using monoclonal anti-histidine antibodies (D).

Amino acid sequences of PCR amplified C1QBP genes from MEAM1/MED whiteflies and *M. persicae* were aligned and compared. C1QBP protein sequence from MEAM1 and MED whiteflies were 100% identical whereas, the C1QBP protein of *M. persicae* was less than 45% identical to that of the whiteflies. Phylogenetic analysis of amino acid sequences of C1QBP of different hemipteran insect vectors also clustered the aphid sequences distinct from all other hemipterans (Figure 6) indicating lesser homology with other hemipteran C1QBP sequences. However, homology modelling of the C1QBP proteins revealed high similarity in the predicted 3D structures of the two proteins (7 A-C) which probably contributes to their interaction with W-CP and Y-CP.

**Figure 6.**
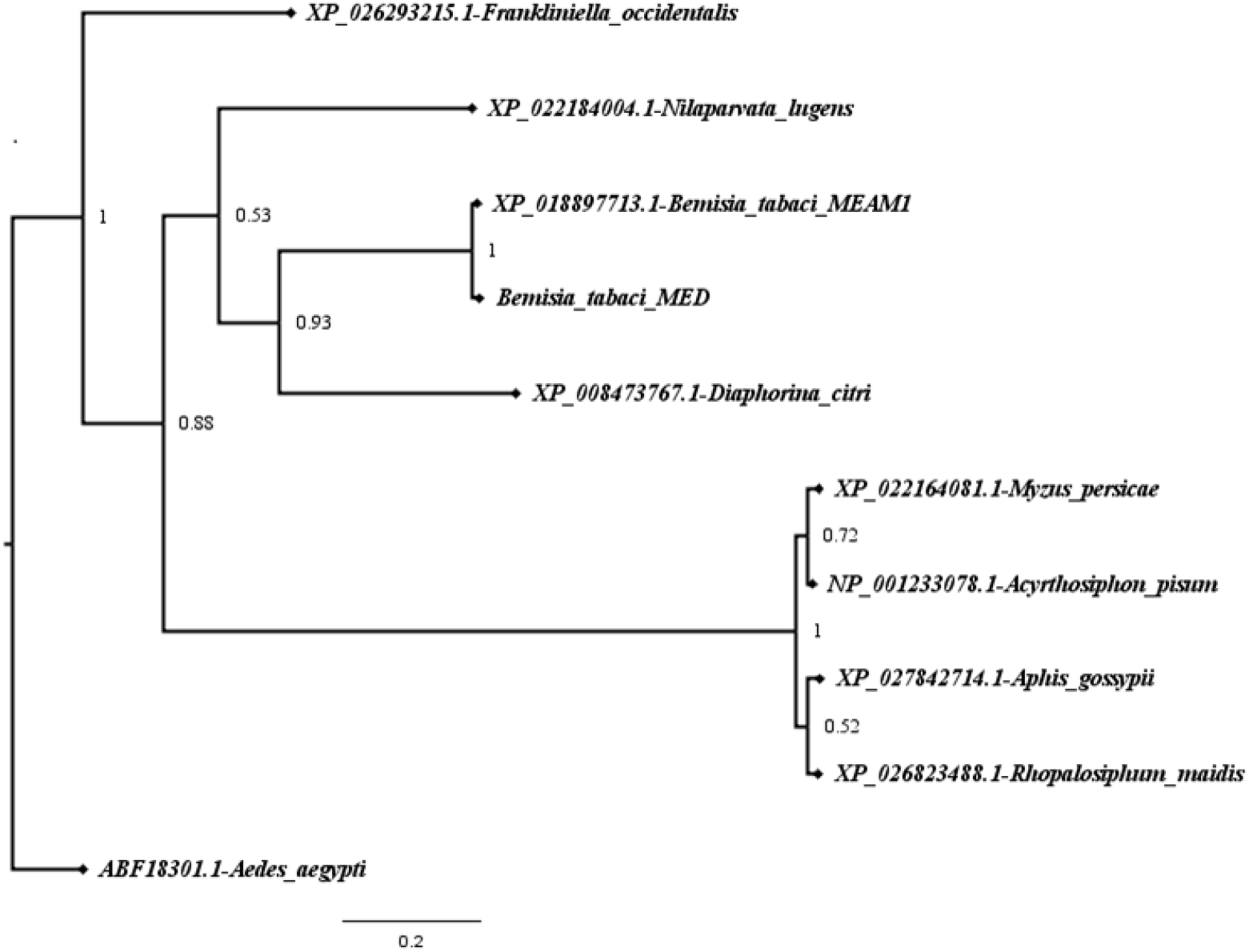
Phylogeny of amino acid sequences of C1QBP receptor proteins of hemipteran insect vectors of plant viruses with *Aedes aegyptii* (yellow fever mosquito) used as an out group.

**Figure 7.**
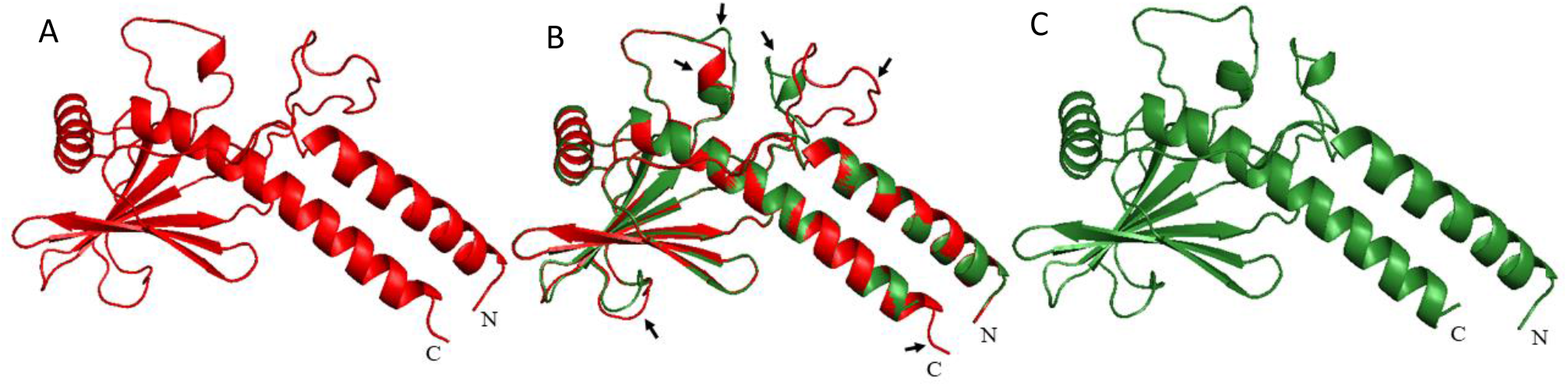
Homology model structures of C1QBP proteins of *B. tabaci* (A), *M. persicae* (C), and their superimposed structures (*B. tabaci*-red and *M. persicae* - green) for comparison (B) with the arrows indicating differences between the two structures.

### Localization of PeWBVYV in whitefly tissues

PeWBVYV was detected as aggregates on the midguts of the MEAM1 whiteflies, concentrated on the ascending and descending loops of the midgut with comparatively lower presence in the filter chamber (Figure 8 A-F). PeWBVYV was also detected from the hindguts of the whitefly. However, PeWBVYV could not be detected in the salivary glands and is possibly due to lower virus titers inside the salivary glands.

**Figure 8.**
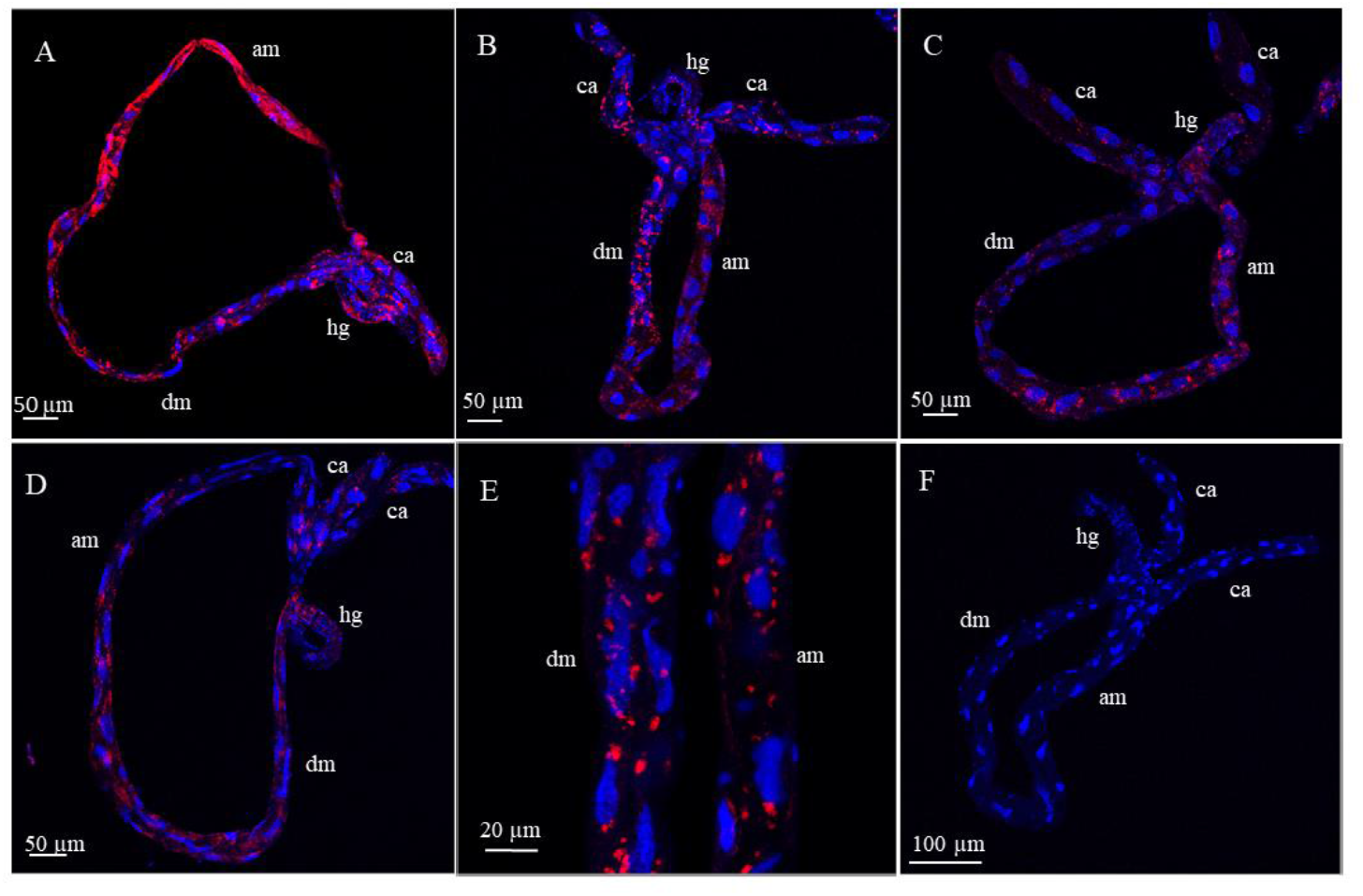
Localization of PeWBVYV on midguts of viruliferous MEAM1 whiteflies at 20x (A-D) and 60x (E) magnification by FISH. Midgut dissected from non-viruliferous whitefly (F) as negative control. am, dm, ca and hg imply ascending midgut, descending midgut, caeca and hindgut, respectively.

## Discussion

Transmission of begomoviruses (persistently transmitted) by *Bemisia tabaci* is hypothesized to be influenced by interaction of the virus capsid proteins with the GroEL chaperone secreted by *Hamiltonella defensa*, a secondary symbiont of the *Bemisia tabaci* which stabilizes the virions under hostile environment (33, 34). The presence and absence of *Hamiltonella* in the MEAM1 and MED whitefly species of Israel, respectively has been correlated with the poor transmission efficiency of TYLCV by the latter (35). In the current study we investigated the transmission of PeWBVYV by MED whiteflies, the other predominant species of *B. tabaci* in Israel. PeWBVYV could not be transmitted by MED whiteflies although the presence of the virus in the whitefly was confirmed by PCR after acquisition. The cause for the specific transmission of PeWBVYV only by MEAM1 whiteflies is unknown. In accordance with known interactions between bacterial GroEL and virus capsid proteins, we tested for interaction between RTD proteins of the whitefly transmitted PeWBVYV and the aphid transmitted PeVYV-2 with the *Hamiltonella* GroEL by Y2H assay. However, unlike TYLCV, the RTD proteins of the poleroviruses screened did not interact with the *Hamiltonella* GroEL. This indicates limited role of *Hamiltonella* GroEL in PeWBVYV transmission and also reasons for non-transmission of PeWBVYV by MED whiteflies remain unknown. Interestingly, we could detect PeVYV-2 and PeWBVYV in the hemolymph of the non-transmitting MEAM1 and MED whiteflies, respectively. In contrast, PeWBVYV could not be detected in the aphid hemolymph indicating its inability to translocate across the midgut barrier and possibly explains its non-transmissibility by aphids. Translocation of the non-transmitted virus across the midgut barrier of the non-transmitter whiteflies but not aphids indicates the semi-specific interaction of the virus particles with the midgut barrier. Moreover, we show that the relative amounts of PeVYV-2 and PeWBVYV in the hemolymph of the non-transmitting MEAM1 and MED, respectively was much lower compared to relative amounts of PeWBVYV in the hemolymph of MEAM1. Mere ability of persistent viruses to cross the midgut barrier and presence in the hemolymph of their insect vectors does not guarantee its transmission (20, 35, 40, 41) but virus quantities above a threshold level within the hemolymph is crucial for its successful transmission (17, 24, 42). Thus, the non-transmission of PeWBVYV by MED whiteflies is possibly due to lower translocation of PeWBVYV across the midgut barrier of the MED whiteflies.

The structural proteins of poleroviruses, primarily the minor capsid protein (RTD) are key determinants of transmission by aphids (21, 26, 27). The amino acid sequence of the RTD of the whitefly-transmitted PeWBVYV differed from that of the known complete RTDs of aphid-transmitted PeVYVs by greater than 10% (3), suggesting a possible role for its altered whitefly transmission. In this study, we screened the minor capsid protein of PeWBVYV against the whitefly cDNA library using a yeast two-hybrid approach. Although we could not identify any whitefly protein exclusively binding with the RTD, however, a receptor protein, complement component 1 Q sub-component binding protein (C1QBP) of the whitefly and aphid was identified to interact with the major coat protein (CP) of PeWBVYV and PeVYV-2. C1QBP is a hexameric pattern recognition glycoprotein localized from diverse cellular components including mitochondria, nucleus, cytoplasm, plasma membrane and even extracellular matrix (43, 44) with ability to identify structures and ligands on microbial surfaces or apoptotic cells (45) and is an important pathogen recognition receptor (43, 44). C1QBP functions as a receptor for bacterial and viral pathogens to facilitate cell adhesion and cellular entry (46–49). C1QBP interactions with viral proteins aid transcriptional activation of viral protein (50, 51), increase viral replication and virion production (52, 53), viral RNA transport (54) or suppress host immune system to promote infection (55, 56). For the first time, interactions of a plant infecting virus with the C1QBP of its insect vector is being reported in this study. Although the function of C1QBP in the transmission of poleroviruses is not known, yet exclusive interaction of the whitefly C1QBP with both PeVYV-2 and PeWBVYV poleroviruses (but not with the whitefly transmitted TYLCV) indicate a specific role in polerovirus transmission. Moreover, the C1QBP protein of *M. persicae*, which is only less than 45 % identical to that of its whitefly homolog interacted strongly with PeVYV-2 and also weakly with PeWBVYV. This further indicates a general receptor-like role for C1QBP for pepper infecting poleroviruses. Recognition of poleroviruses by intestinal receptors although are semi-specific in nature but still are crucial for receptor-mediated endocytotic uptake of virions (7, 29, 30) and also determines the site of virus uptake in the aphid gut (11). The binding of the whitefly C1QBP to both PeWBVYV and PeVYV-2 though fails to reason the transmission of PeWBVYV by whiteflies and not by aphids, still is an important addition as a new candidate receptor for polerovirus with putative role in transmission. Nevertheless, the strong interaction of PeVYV-2 with both the whitefly and aphid C1QBP receptors could lead to competition between the two viruses for transmission by its insect vector. Direct competition between luteoviruses for binding to receptor sites within the aphid vector can significantly affect the transmission efficiency of the competing viruses (15, 57). In our recent study we show that PeWBVYV and PeVYV-2 compete inside their insect vectors with PeWBVYV being the dominant competitor (25). Prior presence of PeWBVYV inside aphids impaired the amounts of PeVYV-2 acquired by it resulting in reduced transmission. However, prior presence of PeVYV-2 inside whiteflies had minimal impact on amounts of PeWBVYV acquired and transmission efficiency. The binding of PeVYV-2 and PeWBVYV to the common C1QBP receptor protein could be a possible reason for such competitive interactions within the insect vectors. Moreover, the detection of PeVYV-2 in the hemolymph of whiteflies is also indicative of binding of the PeVYV-2 virions to the whitefly C1QBP receptor for translocation across the midgut barrier. In contrast, the non-detectability of PeWBVYV in the aphid hemolymph could possibly be due to its lower affinity to the C1QBP receptor of *M. persicae*, as shown in this study. However, it still remains unknown why PeVYV-2 acquisition and transmission by aphids is affected more in presence of PeWBVYV compared to PeWBVYV in MEAM1 whiteflies containing PeVYV-2.

## Conclusions

In this study, we demonstrated that MED species of *B. tabaci* from Israel cannot transmit the newly identified MEAM1-transmitted PeWBVYV polerovirus. The inability of MED to transmit this new virus is not due to absence of the GroEL protein of the *Hamiltonella* symbiont, which has been previously implicated in begomovirus transmission. We show in our study that PeWBVYV crosses the midgut barrier of the MED whiteflies to reach the hemolymph but in much lesser amounts than that of the MEAM1 whiteflies. PeVYV-2, which is transmitted exclusively by aphids was also found to cross the midgut barrier of the whiteflies but in minimal amounts. Moreover, we report a new candidate receptor, C1QBP, from the whitefly and aphids which specifically bind to the major coat protein of poleroviruses. C1QBP is a well-known receptor of pathogenic bacteria and viruses, but this is the first time we report its interactions with a plant virus.

## Author Contributions

Conceptualization, Investigation, Data curation and Formal analysis-M.G. and S.G; Project administration, M.G; Supervision-M.G; Investigation-SG and VHB, Writing Original draft-S.G; review and editing-S.G, VHB and M.G.

## Funding

This project has received funding from the Chief Scientist of the Ministry of Agriculture in Israel grant no 20-02-0102.

## Acknowledgments

The authors wish to thank all members of Ghanim Lab for their support, Amit Gal-On (Plant Pathology, Volcani Center, ARO) for the vectors used for Y2H analysis.

## Conflicts of Interest

The authors declare no conflict of interest. The funders had no role in the design of the study; in the collection, analyses, or interpretation of data; in the writing of the manuscript, or in the decision to publish the results.

## Notes

### Competing Interest Statement

The authors have declared no competing interest.

## References

1. Power AG. 2000. Insect transmission of plant viruses: a constraint on virus variability. Curr Opin Plant Biol 3:336–340.

2. Lefeuvre P, Martin DP, Elena SF, Shepherd DN, Roumagnac P, Varsani A. 2019. Evolution and ecology of plant viruses. Nat Rev Microbiol 17:632–644.

3. Ghosh S, Kanakala S, Lebedev G, Kontsedalov S, Silverman D, Alon T, Mor N, Sela N, Luria N, Dombrovsky A, Mawassi M, Haviv S, Czosnek H, Ghanim M. 2019. Transmission of a new polerovirus infecting pepper by the whitefly Bemisia tabaci. J Virol 93:e00488–19.

4. Costa TM, Inoue-Nagata AK, Vidal AH, Ribeiro SG, Nagata T. 2020. The recombinant isolate of cucurbit aphid-;borne yellows virus from Brazil is a polerovirus transmitted by whiteflies. Plant Pathol.

5. Gilbertson RL, Batuman O, Webster CG, Adkins S. 2015. Role of the insect supervectors Bemisia tabaci and Frankliniella occidentalis in the emergence and global spread of plant viruses. Annu Rev Virol 2:67–93.

6. Gray S, Gildow FE. 2003. Luteovirus-aphid interactions. Annu Rev Phytopathol 41:539–566.

7. Gildow FE. 1993. Evidence for receptor-mediated endocytosis regulating Luteovirus acquisition by aphids. Phytopathology 83:270.

8. Brault V, Herrbach É, Reinbold C. 2007. Electron microscopy studies on luteovirid transmission by aphids. Micron 38:302–312.

9. Ghanim M, Morin S, Czosnek H. 2001. Rate of Tomato yellow leaf curl virus translocation in the circulative transmission pathway of its vector, the whitefly Bemisia tabaci. Phytopathology 91:188–196.

10. Uchibori M, Hirata A, Suzuki M, Ugaki M. 2013. Tomato yellow leaf curl virus accumulates in vesicle-like structures in descending and ascending midgut epithelial cells of the vector whitefly, Bemisia tabaci, but not in those of nonvector whitefly Trialeurodes vaporariorum. J Gen Plant Pathol 79:115–122.

11. Gray S, Gildow FE. 2003. Luteovirus-aphid interactions. Annu Rev Phytopathol 41:539–566.

12. Brault V, Herrbach É, Reinbold C. 2007. Electron microscopy studies on luteovirid transmission by aphids. Micron 38:302–312.

13. Pan LL, Chen QF, Zhao JJ, Guo T, Wang XW, Hariton-Shalev A, Czosnek H, Liu SS. 2017. Clathrin-mediated endocytosis is involved in Tomato yellow leaf curl virus transport across the midgut barrier of its whitefly vector. Virology 502:152–159.

14. Xia WQ, Liang Y, Chi Y, Pan LL, Zhao J, Liu SS, Wang XW. 2018. Intracellular trafficking of begomoviruses in the midgut cells of their insect vector. PLoS Pathog 14.

15. Gildow FE, Rochow WF. 1980. Role of accessory salivary glands in aphid transmission of barley yellow dwarf virus. Virology 104:97–108.

16. Czosnek H, Ghanim M, Ghanim M. 2002. The circulative pathway of begomoviruses in the whitefly vector Bemisia tabaci - Insights from studies with Tomato yellow leaf curl virus. Ann Appl Biol 140:215–231.

17. Pan L, Chen Q, Guo T, Wang X, Li P, Wang X, Liu S. 2018. Differential efficiency of a begomovirus to cross the midgut of different species of whiteflies results in variation of virus transmission by the vectors. Sci China Life Sci 61:1254–1265.

18. Gildow FE, Gray SM. 1993. The aphid salivary gland basal lamina as a selective barrier associated with vector-specific transmission of barley yellow dwarf luteoviruses. Phytopathology 83:1293–1302.

19. Ohnishi J, Kitamura T, Terami F, Honda KI. 2009. A selective barrier in the midgut epithelial cell membrane of the nonvector whitefly Trialeurodes vaporariorum to Tomato yellow leaf curl virus uptake. J Gen Plant Pathol 75:131–139.

20. Wei J, Zhao J-J, Zhang T, Li F-F, Ghanim M, Zhou X-P, Ye G-Y, Liu S-S, Wang X-W. 2014. Specific Cells in the Primary Salivary Glands of the Whitefly Bemisia tabaci Control Retention and Transmission of Begomoviruses. J Virol 88:13460–13468.

21. Bruyere A, Brault V, Ziegler-Graff V, Simonis M-T, Van Den Heuvel J, Richards K, Guilley H, Jonard G, Herrbach E. 1997. Effects of mutations in the beet western yellows virus readthrough protein on its expression and packaging and on virus accumulation, symptoms, and aphid transmission. Virology 230:323–334.

22. Brault V, Périgon S, Reinbold C, Erdinger M, Scheidecker D, Herrbach E, Richards K, Ziegler-Graff V. 2005. The polerovirus minor capsid protein determines vector specificity and intestinal tropism in the aphid. J Virol 79:9685–9693.

23. Noris E, Vaira AM, Caciagli P, Masenga V, Gronenborn B, Accotto GP. 1998. Amino Acids in the Capsid Protein of Tomato Yellow Leaf Curl Virus That Are Crucial for Systemic Infection, Particle Formation, and Insect Transmission. J Virol 72:10050–10057.

24. Guo T, Zhao J, Pan LL, Geng L, Lei T, Wang XW, Liu SS. 2018. The level of midgut penetration of two begomoviruses affects their acquisition and transmission by two species of Bemisia tabaci. Virology 515:66–73.

25. Bello VH, Ghosh S, Krause-Sakate R, Ghanim M. 2020. Competitive Interactions Between Whitefly and Aphid Transmitted Poleroviruses within the Plant Host and the Insect Vectors. Phytopathology®.

26. Brault V, Mutterer J, Scheidecker D, Simonis MT, Herrbach E, Richards K, Ziegler-Graff V. 2000. Effects of point mutations in the readthrough domain of the beet western yellows virus minor capsid protein on virus accumulation in planta and on transmission by aphids. J Virol 74:1140–1148.

27. Reinbold C, Gildow FE, Herrbach E, Ziegler-Graff V, Goncalves MC, Van Den Heuvel J, Brault V. 2001. Studies on the role of the minor capsid protein in transport of Beet western yellows virus through Myzus persicae. J Gen Virol 82:1995–2007.

28. Chay CA, Gunasinge UB, Dinesh-Kumar SP, Miller WA, Gray SM. 1996. Aphid transmission and systemic plant infection determinants of barley yellow dwarf luteovirus-PAV are contained in the coat protein readthrough domain and 17-kDa protein, respectively. Virology 219:57–65.

29. Linz LB, Liu S, Chougule NP, Bonning BC. 2015. In Vitro Evidence Supports Membrane Alanyl Aminopeptidase N as a Receptor for a Plant Virus in the Pea Aphid Vector. J Virol 89:11203–11212.

30. Mulot M, Monsion B, Boissinot S, Rastegar M, Meyer S, Bochet N, Brault V. 2018. Transmission of Turnip yellows virus by Myzus persicae is reduced by feeding aphids on double-stranded RNA targeting the ephrin receptor protein. Front Microbiol 9.

31. Wang X, Zhou G. 2003. Identification of a protein associated with circulative transmission of Barley yellow dwarf virus from cereal aphids, Schizaphis graminum and Sitobion avenae. Chinese Sci Bull 48:2083–2087.

32. Van den Heuvel JF, Bruyere A, Hogenhout SA, Ziegler-Graff V, Brault V, Verbeek M, Van Der Wilk F, Richards K. 1997. The N-terminal region of the luteovirus readthrough domain determines virus binding to Buchnera GroEL and is essential for virus persistence in the aphid. J Virol 71:7258–7265.

33. Morin S, Ghanim M, Zeidan M, Czosnek H, Verbeek M, van den Heuvel JFJM. 1999. A GroEL Homologue from Endosymbiotic Bacteria of the WhiteflyBemisia tabaciIs Implicated in the Circulative Transmission of Tomato Yellow Leaf Curl Virus. Virology 256:75–84.

34. Morin S, Ghanim M, Sobol I, Czosnek H. 2000. The GroEL protein of the whitefly Bemisia tabaci interacts with the coat protein of transmissible and nontransmissible begomoviruses in the yeast two-hybrid system. Virology 276:404–416.

35. Gottlieb Y, Zchori-Fein E, Mozes-Daube N, Kontsedalov S, Skaljac M, Brumin M, Sobol I, Czosnek H, Vavre F, Fleury F. 2010. The transmission efficiency of tomato yellow leaf curl virus by the whitefly Bemisia tabaci is correlated with the presence of a specific symbiotic bacterium species. J Virol 84:9310–9317.

36. El-Gebali S, Mistry J, Bateman A, Eddy SR, Luciani A, Potter SC, Qureshi M, Richardson LJ, Salazar GA, Smart A. 2018. The Pfam protein families database in 2019. Nucleic Acids Res 47:D427–D432.

37. Marchler-Bauer A, Derbyshire MK, Gonzales NR, Lu S, Chitsaz F, Geer LY, Geer RC, He J, Gwadz M, Hurwitz DI. 2014. CDD: NCBI’s conserved domain database. Nucleic Acids Res 43:D222–D226.

38. Kumar S, Stecher G, Tamura K. 2016. MEGA7: molecular evolutionary genetics analysis version 7.0 for bigger datasets. Mol Biol Evol 33:1870–1874.

39. Miller MA, Pfeiffer W, Schwartz T. 2010. Creating the CIPRES Science Gateway for inference of large phylogenetic trees, p. 1–8. In Gateway Computing Environments Workshop (GCE), 2010. Ieee.

40. Rochow WF, Pang E. 1961. Aphids can acquire strains of barley yellow dwarf virus they do not transmit. Virology.

41. Caciagli P, Medina Piles V, Marian D, Vecchiati M, Masenga V, Mason G, Falcioni T, Noris E. 2009. Virion Stability Is Important for the Circulative Transmission of Tomato Yellow Leaf Curl Sardinia Virus by Bemisia tabaci, but Virion Access to Salivary Glands Does Not Guarantee Transmissibility. J Virol 83:5784–5795.

42. Pan LL, Chi Y, Liu C, Fan YY, Liu SS. 2020. Mutations in the coat protein of a begomovirus result in altered transmission by different species of whitefly vectors. Virus Evol 6.

43. Peerschke EIB, Ghebrehiwet B. 2007. The contribution of gC1qR/p33 in infection and inflammation. Immunobiology 212:333–342.

44. Pednekar L, Valentino A, Ji Y, Tumma N, Valentino C, Kadoor A, Hosszu KK, Ramadass M, Kew RR, Kishore U. 2016. Identification of the gC1qR sites for the HIV-1 viral envelope protein gp41 and the HCV core protein: Implications in viral-specific pathogenesis and therapy. Mol Immunol 74:18–26.

45. Kouser L, Madhukaran SP, Shastri A, Saraon A, Ferluga J, Al-Mozaini M, Kishore U. 2015. Emerging and novel functions of complement protein C1q. Front Immunol 6:317.

46. Braun L, Ghebrehiwet B, Cossart P. 2000. gC1q-;R/p32, a C1q-;binding protein, is a receptor for the InlB invasion protein of Listeria monocytogenes. EMBO J 19:1458–1466.

47. Nguyen T, Ghebrehiwet B, Peerschke EIB. 2000. Staphylococcus aureus protein A recognizes platelet gC1qR/p33: a novel mechanism for staphylococcal interactions with platelets. Infect Immun 68:2061–2068.

48. Ghebrehiwet B, Tantral L, Titmus MA, Panessa-Warren BJ, Tortora GT, Wong SS, Warren JB. 2007. The exosporium of B. cereus contains a binding site for gC1qR/p33: implication in spore attachment and/or entry, p. 181–197. In Current Topics in Innate Immunity. Springer.

49. Choi Y, Kwon Y-C, Kim S-I, Park J-M, Lee K-H, Ahn B-Y. 2008. A hantavirus causing hemorrhagic fever with renal syndrome requires gC1qR/p32 for efficient cell binding and infection. Virology 381:178–183.

50. Wang Y, Finan JE, Middeldorp JM, Hayward SD. 1997. P32/TAP, a cellular protein that interacts with EBNA-1 of Epstein–Barr virus. Virology 236:18–29.

51. Yu L, Loewenstein PM, Zhang Z, Green M. 1995. In vitro interaction of the human immunodeficiency virus type 1 Tat transactivator and the general transcription factor TFIIB with the cellular protein TAP. J Virol 69:3017–3023.

52. Mohan KVK, Ghebrehiwet B, Atreya CD. 2002. The N-terminal conserved domain of rubella virus capsid interacts with the C-terminal region of cellular p32 and overexpression of p32 enhances the viral infectivity. Virus Res 85:151–161.

53. Hu M, Li H-M, Bogoyevitch MA, Jans DA. 2017. Mitochondrial protein p32/HAPB1/gC1qR/C1qbp is required for efficient respiratory syncytial virus production. Biochem Biophys Res Commun 489:460–465.

54. Luo Y, Yu H, Peterlin BM. 1994. Cellular protein modulates effects of human immunodeficiency virus type 1 Rev. J Virol 68:3850–3856.

55. Kittlesen DJ, Chianese-Bullock KA, Yao ZQ, Braciale TJ, Hahn YS. 2000. Interaction between complement receptor gC1qR and hepatitis C virus core protein inhibits T-lymphocyte proliferation. J Clin Invest 106:1239–1249.

56. Yao ZQ, Eisen-Vandervelde A, Waggoner SN, Cale EM, Hahn YS. 2004. Direct binding of hepatitis C virus core to gC1qR on CD4+ and CD8+ T cells leads to impaired activation of Lck and Akt. J Virol 78:6409–6419.

57. Rochow, W.F.; Muller, I.; Gildow FE. 1983. Interference between two luteoviruses in an aphid: lack of reciprocal competition. Phytopathology 73:919–922.

